# Interfering the mating of *Chilo suppressalis* (Walker): A new role of sex pheromones ((Z)-11-Octadecen-1-ol and (Z)-13-Octadecen-1-ol) from *Cnaphalocrocis medinalis* Guenée

**DOI:** 10.1101/2020.01.03.893339

**Authors:** Yu-yong Liang, Mei Luo, Xiao-gang Fu, Li-xia Zheng, Hong-yi Wei

**Author notes:** Corresponding author. E-mail address (H-Y Wei).

## Abstract

The rice stem borer, *Chilo suppressalis* (Walker), and the rice leaf folder, *Cnaphalocrocis medinalis* Guenée are two of the most destructive lepidopteran pests in rice. Since these two lepidopteran insects show occurrence laps in rice paddy fields, farmers prefer to set the pheromone-baited traps of *C. suppressalis* accompany with the pheromone traps of *C. medinalis* in the rice fields for convenient observation. However, our field observation demonstrated that no male of the rice stem borer was captured in the traps baited with commercial *C. suppressalis* sex pheromone (CCS) combined with commercial *C. medinalis* sex pheromone (CCM). To confirm the *C. medinalis* sex pheromone component(s) to interfere with the attraction of males of the rice stem borers to their conspecific female moths, the single component of *C. medinalis* sex pheromone combined with CCS in traps were tested in the laboratory and in the rice paddy field. The results revealed the two alcohol components in CM, i.e., (Z)-11-octadecen-1-ol (Z11-18:OH) and (Z)-13-octadecen-1-ol (Z13-18:OH) may cause a significant reduction in the captures by CCS to *C. suppressalis* males. We recommend against using these sex pheromones together in the field and suggested that Z11-18:OH and Z13-18:OH could be potential inhibitors or antagonists of *C. suppressalis* sex pheromone to control the rice stem borer in practice.

## Introduction

The rice stem borer, *Chilo suppressalis* (Walker) and the rice leaf folder, *Cnaphalocrocis medinalis* Guenée, are two of the most harmful lepidopteran rice pests throughout China and other Asian countries (Su et al. 2014, Luo et al. 2019). In China, these two pests caused wide and extensive damages in rice and led to great economic loss in recent decades (Sheng et al. 2003, Liu et al. 2008). Currently, chemical control, such as application of insecticides has been utilized as an efficient method to control these two pests in China (Huang et al. 2011, Zheng et al. 2011). However, overuse of insecticides may cause severe insecticide resistance and serious environmental problems (Chen and Klein 2012, Fu et al. 2018, Pu et al. 2019). Therefore, biological control has been raised to be the one of most important strategies to suppress the outbreaks of these rice pests (Lou et al. 2014).

Sex pheromone application is becoming a valuable and efficient strategy of biological control to suppress lepidopteran pests (Chen et al. 2014). For example, pheromone-baited traps were widely used in male flight monitoring, population forecasting, mass trapping, and mating disruption to monitor and control a lot of lepidopteran pest species (Campion and Nesbitt 1983, Witzgall et al. 2010, Chen et al. 2014). Compared to the conventional methods, the pheromone-based control methods represent the advantages via the species-specificity of pheromones. The high biological activity which means that only relatively small amounts are required, and the negligible toxic effects on plants and animals (Campion and Nesbitt 1983). Pheromones generally consist of various component, which may be isomers with respect to the geometry and position of the unsaturation or they may be structurally related compounds differing in chain length or the nature of the functional group. In most cases, the components are secreted in a very precise ratio (Campion and Nesbitt 1983).

In most lepidopteran insects, sex pheromones are generally secreted by female moths for attracting the males for copulation (Raina 1989). The *C. suppressalis* sex pheromone was first identified by Nesbitt et al. (1975) as a two-component blend of (Z)-11-hexadecenal (Z11-16:Ald) and (Z)-13-octadecenal (Z13-18:Ald). In 1983, an additional compound, (Z)-9-hexadecenal (Z9-16:Ald), was identified (Tatsuki et al. 1983). These three-component blends (Z11-16:Ald, Z13-18:Ald, and Z9-16:Ald) were found at a ratio of 46:6:5, which is used in production of the standard lure in *C. suppressalis* (Tatsuki 1990, Cork 2004). In general, the *C. medinalis* sex pheromone consists of four components, (Z)-11-octadecenal (Z11-18:Ald), (Z)-13-octadecenal (Z13-18:Ald), (Z)-11-octadecen-1-ol (Z11-18:OH) and (Z)-13-octadecen-1-ol (Z13-18:OH) as a mixed ratio of 11:100:24:36 (Kawazu et al. 2000). Since then, the sex pheromones of *C. suppressalis* and *C. medinalis* were widely applied in the rice paddies and has become an ideal way for integrated pest management (IPM) programs (Kawazu et al. 2004, Kawazu et al. 2005, Byers 2007, Litsinger 2009, Cho et al. 2013, Chen et al. 2014).

*Chilo suppressalis* occurs four generations a year in most parts of Jiangxi province, China. Moreover, the emergence period of *C. suppressalis* and *C. medinalis* overlapped extensively. In this case, the two sex pheromones may be used at the same time by the famers to control *C. suppressalis* and *C. medinalis* together. However, previous studies have reported that mixing sex pheromones of two pest species resulted in interference of capture (Haynes et al. 2002, Gemeno et al. 2006). In addition, weather mixing the sex pheromones of *C. suppressalis* and *C. medinalis* is risky in interference of capture is unknown now. Therefore, in this study, we aimed to determine whether mixing the sex pheromones of *C. suppressalis* and *C. medinalis* has negative effects in the IPM. We also want to determine the mechanism if mixing the sex pheromones has negative effects in pest control. This study is very important for guiding farmers in intensive application of various sex pheromones in practice.

## Materials and Methods

### Insects

A laboratory colonized population of *C. suppressalis* was collected from Shang-gao county, Jiangxi, China (E115°04’53.34”, N28°19’10.87”). Naturally overwintering larvae were collected from the paddy field in late March 2012, then were successively reared in cylindrical glass jars (10.0 cm in height and 12.0 cm in diameter) with several rice stems. Cotton stoppers for jars to prevent insects from escaping. The rice stems were changed every two days. Pupae were collected and transferred to 24-well plates for emergence individually. New emerged female and male adults were maintained separately in transparent plastic bags with a 10% sucrose solution. The insects were maintained in a growth room under a 14:10 (L: D) photo regime at 25 ± 1C° and 85 ± 5% RH.

### Chemicals

The commercial sex pheromones of *C. suppressalis* (CCS) and *C. medinalis* (CCM) were purchased from Pherobio Technology Co., Ltd (Beijing, China). The synthetic chemicals Z11-16:Ald, Z13-18:Ald, Z9-16:Ald, Z11-18:Ald, Z11-18:OH and Z13-18:OH with more than 99% purity obtained from Sigma^®^ Chemical Co. (St. Louis, MO, U.S.A.).

### Experimental devices

The trap 1 (Fig. 1a) is custom-made, and it consists of a basin (25.0 cm in diameter and 10.0 cm in deep) and a triangle bracket. Two drainage holes were formed away from the dish bottom 6.0-7.0 cm. The basin was fixed on the bracket, and the height of the basin was always adjusted to 10.0-20.0 cm higher than the rice. 2.0 g washing powder were added to the water in the basin. Subsequently, the basin was filled with water until the drainage holes. The lures were always fixed 0.5-1.0 cm above the water.

**Fig. 1.**
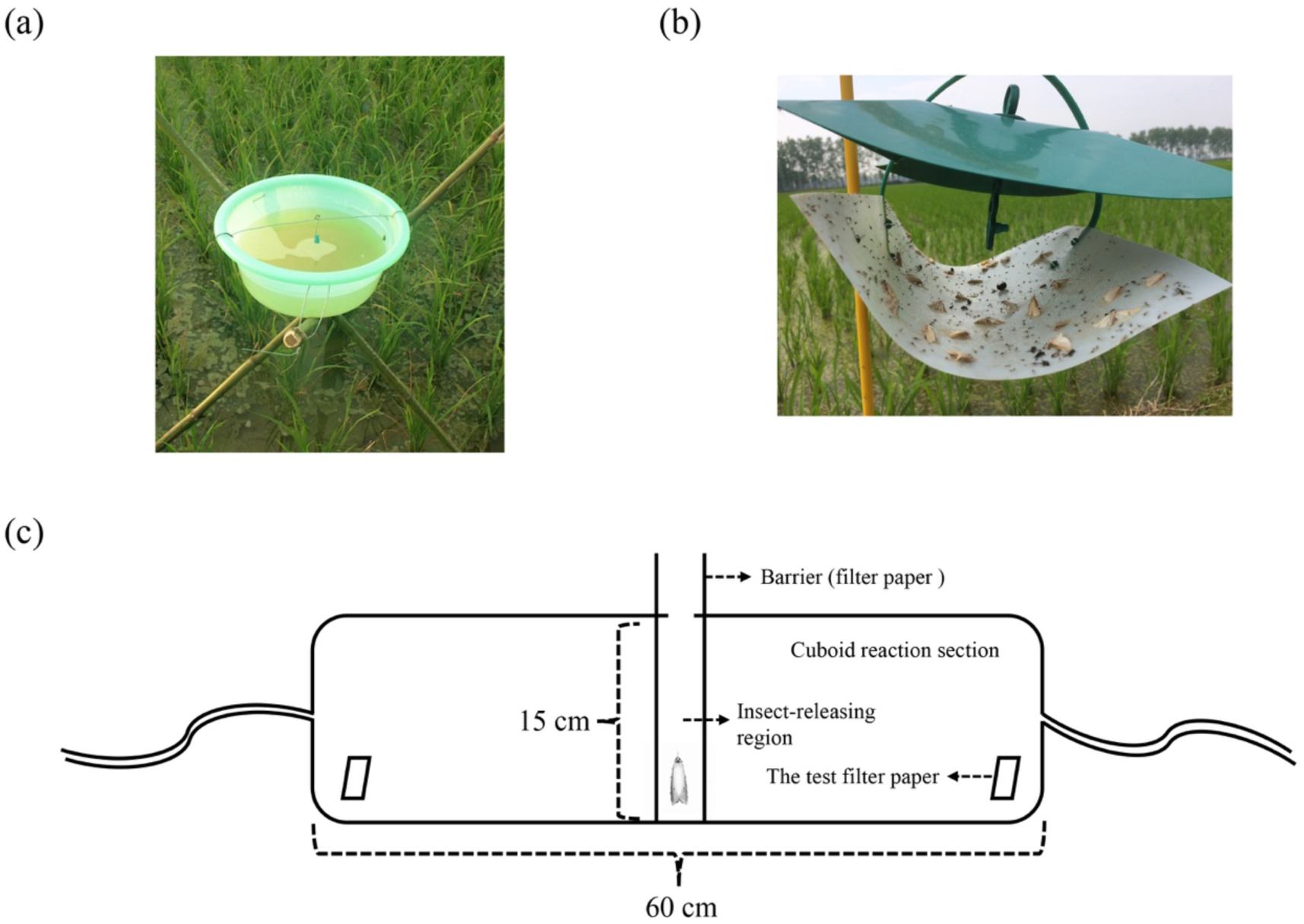
The schematic diagrams of three experimental devices in this study. (a) trap 1, (b) trap 2 and (c) olfactometer.

The trap 2 (Fig. 1b) was purchased from Pherobio Technology Co., Ltd (Beijing, China), which consists of a ship-type trap and a trestle. The ship-type trap consists of a sticky board, a bezel and a jack device for lures.

The olfactometer (Fig. 1c) is custom-made, and it consists of a cuboid reaction section of 60.0 cm long by 15.0 cm inner diameters. The cuboid was averagely divided into three parts, an insect-releasing region of the middle part and two testing regions at the two ends, respectively. The insect-releasing region and testing regions was baffled with filter papers. An exit opening was located on the back of the device. The olfactometer was kept on the table, humidified and purified air at 1.0 L/min on both ends was injected into the device. The lures were made of strips of filter paper (1.0 cm × 5.0 cm), which loaded with either 100 μl of the test stimulus, or redistilled hexane (control).

### Experimental design

To determine whether mixing the sex pheromones of *C. suppressalis* and *C. medinalis* interfere the capture of *C. suppressalis* and *C. medinalis*, we tested the attraction of the traps baited with CCS and CCM to these two pests in the field (Experiment 1) (Table 1). The traps (trap 1) with two sex pheromones and CCS (control) were tested with 3 traps for 3 replicates from July to September 2012. Trap captures were counted every day.

**Table 1.** The experimental design.

| Experiment | Experimental purpose | Experimental device | Time |
| --- | --- | --- | --- |
| Experiment 1 | To determine whether mixing the sex pheromones of *C. suppressalis* and *C. medinalis* interfere the capture of *C. suppressalis* and *C. medinalis* | Trap 1 | July to September 2012 |
| Experiment 2 | To determine the optimal concentration of CS components in behavioral experiments | Wind tunnel | July to August 2013 |
| Experiment 3 | To determine the response of *C. suppressalis* male to sex pheromone components of *C. medinalis* | Olfactometer | July to August 2013 |
| Experiment 4 | To determine the effects of the sex pheromone components of *C. medinalis* on *C. suppressalis* in the field | Trap 2 | June to August 2014 |

To determine the optimal concentration of CS (three components of *C. suppressalis* sex pheromone (Z11-16:Ald, Z13-18:Ald, Z9-16:Ald=46:6:5)) in behavioral experiments, we tested responses of *C. suppressalis* males in the wind tunnel to its sex pheromones at different concentrations (Experiment 2) (Table 1). The wind tunnel was a cylindrical (2.0 m in length and 0.5 m in diameter) aluminum frame with glass walls. The experimental operation of the wind tunnel is the same with it in our another experiment (Luo et al. 2020). The pheromone source with different concentrations (0.1, 1.0, 10.0 μg/μl) of CS were placed in a 4.0 cm^2^ mesh cage and were placed 20.0 cm from the upper air outlet of wind tunnel. One-day-old male adults were placed in a steel screen platform which was placed above the floor of the wind tunnel. Thirty males were tested for each treatment and each male was tested only once. After release, male behavior was observed for 2.0 min. Behaviors recorded included exciting, taking flight from the release cage, oriented flight, contacting the pheromone source.

To determine the response of male *C. suppressalis* to sex pheromone components of *C. medinalis*, we tested the effects of different sex pheromone components of *C. medinalis* to male *C. suppressalis* with a two-way choice bioassay by using a custom-made olfactometer (Experiment 3) (Table 1). For each bioassay, male *C. suppressalis* individually was introduced into the insect-releasing region of the device. The males were released after the barriers (two filter papers) were pulled out. The male *C. suppressalis* initiated flight and migrated to one of the testing ends (set over half of the choice chambers) and the choice was recorded after it remained for at least 60.0 s in that testing end. If the male did not make a choice within 5.0 min after being released into the olfactometer, it was considered as a non-responder. We rotated the olfactometer 180° to randomize any positional effects after five males have been tested. We cleared the olfactometer with 95% ethyl alcohol after ten males have been tested. The test filter papers were used as the lures in this study, which containing different sex pheromone components. The lures were CCM; CCM+CCS; Z11-18:Ald; Z13-18:OH; Z11-18:OH; four-component blend of CCM, at a ratio of 100:36:11:24, of the Z13-18:Ald, Z13-18:OH, Z11-18:Ald and Z11-18:OH (CM); the components of CCS and CCM, at a ratio of 140:117:15:36:11:24, of the Z11-16:Ald, Z13-18:Ald, Z9-16:Ald, Z13-18:OH, Z11-18:Ald, Z11-18:OH (CS+CM). Thirty males were tested in each treatment.

To determine the effects of the sex pheromone components of *C. medinalis* on *C. suppressalis* in the paddy field, we tested the attraction of male *C. suppressalis* to the traps baited with a combination of CCS and the components of CCM by the trap 2 (Experiment 4) (Table 1). Combined Z11-18:Ald, Z13-18:OH and Z11-18:OH with CCS respectively were the treatments and CCS alone was the control treatment. Each treatment was tested with 3 traps for 3 replicates from June to August 2014. Trap captures were also counted every day.

### Data analysis

We compared the difference of capture of *C. suppressalis* males between treatments and control treatment with a *t*-test by SPSS 24.0 (SPSS Inc., Chicago, IL). Differences in olfactometer experiments were analyzed by a *Chi*-square test. The percentage data were arcsine-square-root transformed and statistically evaluated by one-way ANOVA followed by Duncan’s new multiple range test. Significance level was set at 0.05.

## Results

### Mixing the sex pheromones of *C. suppressalis* and *C. medinalis* interfered the capture of *C. suppressalis*

During the experimental period (July to September 2012), we found that no *C. suppressalis* male and ten *C. medinalis* males were caught in the traps baited with combination of CCS and CCM. However, the total captures by CCS were 254 and the average trap caught by CCS during the trial was 2.41 males per trap per day, which were extremely significant higher than those in treatment of combining CCS and CCM (*t* _(76)_ = 6.045, *P* < 0.001) (Fig. 2). The peak point of male *C. suppressalis* captured by its sex pheromone (CCS) appeared on August 29th, 2012. While, there was zero *C. suppressalis* caught by the combination of CCS and CCM during the whole period. This result suggested that mixing the sex pheromones of *C. suppressalis* and *C. medinalis* interfered the capture of *C. suppressalis* in the field.

**Fig. 2.**
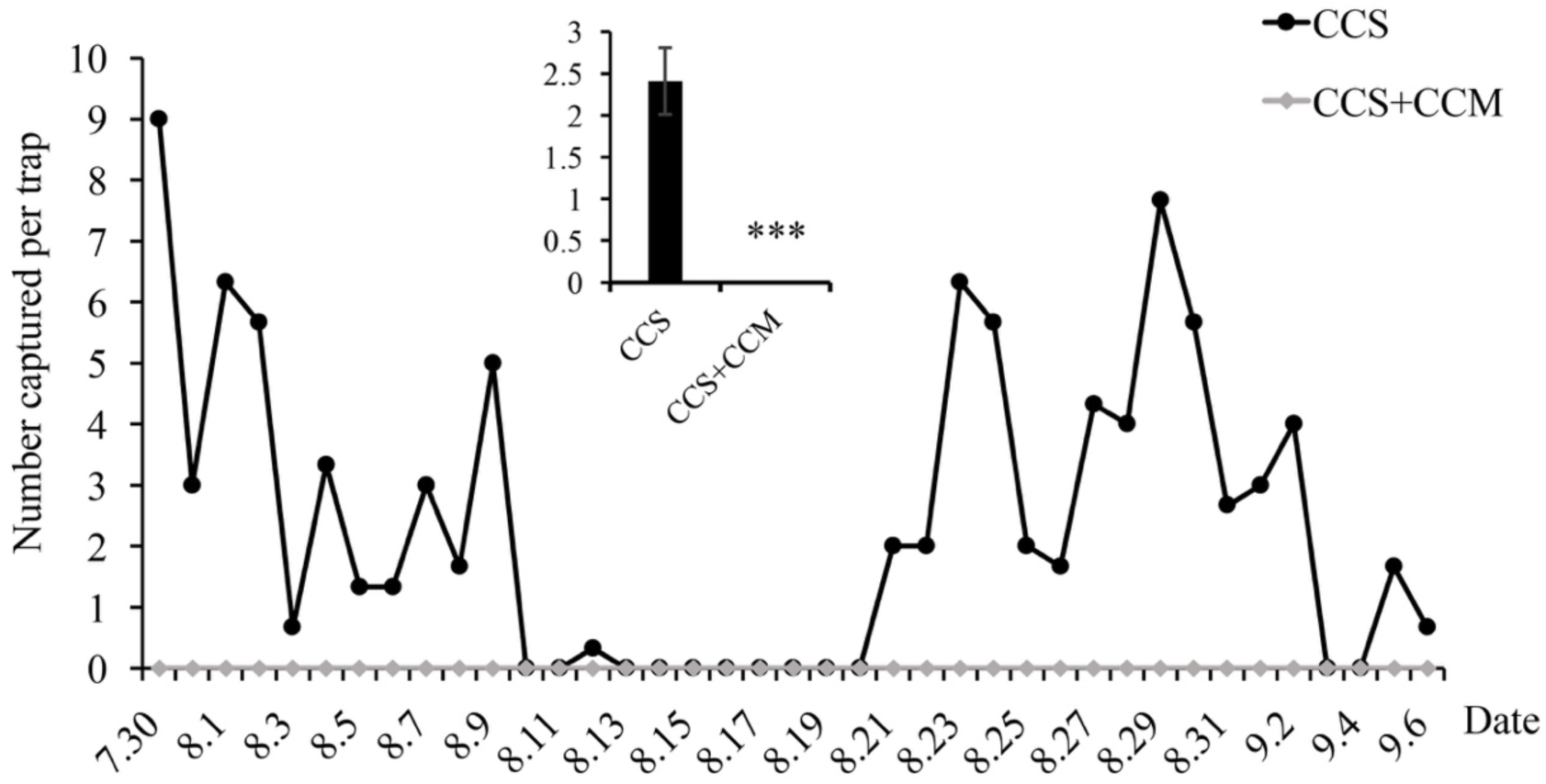
The number of *C. suppressalis* males captured by commercial *C. suppressalis* sex pheromone alone (CCS) (black) and combined with commercial *C. medinalis* sex pheromone (CCS+CCM) (gray) in Jiangxi, China, from July to September 2012. We compared the difference of capture of *C. suppressalis* males between treatments and control treatment with a *t*-test by SPSS 12.0. ***, *P* < 0.001.

### Z11-18:OH and Z13-18:OH are two major components of CCM that inhibited the attraction of male *C. suppressalis* in olfactory behavior

To determine the optimal concentration of CS components in behavioral experiments, we tested the response of *C. suppressalis* males to CCS and three concentrations of CS (Z11-16Ald: Z13-18Ald: Z9-16Ald = 140:17:15) in the wind tunnel experiment (Table 2). The result showed that the CCS gave the highest percentage of *C. suppressalis* males exciting, taking flight, orienting flight and contacting source among the five treatments. Among the three concentrations of the three components (Z11-16Ald: Z13-18Ald: Z9-16Ald = 140:17:15) of CS, *C. suppressalis* males showed high levels of exciting, taking flight, orienting flight and contacting source at the concentration of 1.0 μg/μl. In addition, there was no difference observed in the four behaviors in the concentrations of 0.1, 10.0 μg/μl and control. Therefore, 1.0 μg/μl was used as the optimal concentration of CS in the further behavioral experiments.

**Table 2.** The different behavioral responses of *C. suppressalis* males in the wind tunnel to the sex pheromones at different concentrations

| Treatments | Response (%) |  |  |  |
| --- | --- | --- | --- | --- |
|  | Exciting | Taking flight | Orienting flight | Contacting source |
| CCS | 53.3 ± 5.8a | 36.7 ± 11.5a | 23.3 ± 5.8a | 13.3 ± 5.8a |
| CS (0.1 µg/µl) | 16.7 ± 5.8c | 10.0 ± 0.0b | 6.7 ± 5.8b | 6.7 ± 5.8b |
| CS (1 µg/µl) | 36.7 ± 5.8b | 20.0 ± 0.0b | 10.0 ± 10.0b | 6.7 ± 5.8b |
| CS (10 µg/µl) | 20.0 ± 0.0c | 10.0 ± 10.0b | 0.0 ± 0.0b | 0.0 ± 0.0b |
| CK | 20.0 ± 10.0c | 13.3 ± 5.8b | 0.0 ± 0.0b | 0.0 ± 0.0b |

Data in the table are means ± *SE*, and those in the same column followed by different letters are significantly different ( $P < 0.05$ , ANOVA followed by Duncan's new multiple range test). CCS means commercial *C. suppressalis* sex pheromone. CS means the three components of *C. suppressalis* sex pheromone (Z11-16:Ald, Z13-18:Ald, Z9-16:Ald=46:6:5). The control (CK) is redistilled hexane. $N = 30$ .

To determine the response of *C. suppressalis* male to sex pheromone components of *C. medinalis*, we tested the attractiveness of the seven lures to *C. suppressalis* males by a custom-made olfactometer (Fig. 3). We found that the lures of Z11-18:Ald (*χ*^*2*^ *=* 1.067, *P* = 0.302), Z13-18:OH (*χ*^*2*^ *=* 1.067, *P* = 0.302) and CCS+CCM (*χ*^*2*^ *=* 1.067, *P* = 0.302) have no significant effects on responses of *C. suppressalis* males. However, the males were predominantly attracted to the control than to Z11-18:OH (*χ*^*2*^ *=* 32.267, *P* < 0.001), CM (*χ*^*2*^ *=* 17.067, *P* < 0.001), CS+CM (*χ*^*2*^ *=* 38.400, *P* < 0.001) and CCM (*χ*^*2*^ *=* 13.067, *P* < 0.001) treatments. Which suggested that Z11-18:OH is the major component of *C. medinalis* to inhibit responses of *C. suppressalis* males.

**Fig. 3.**
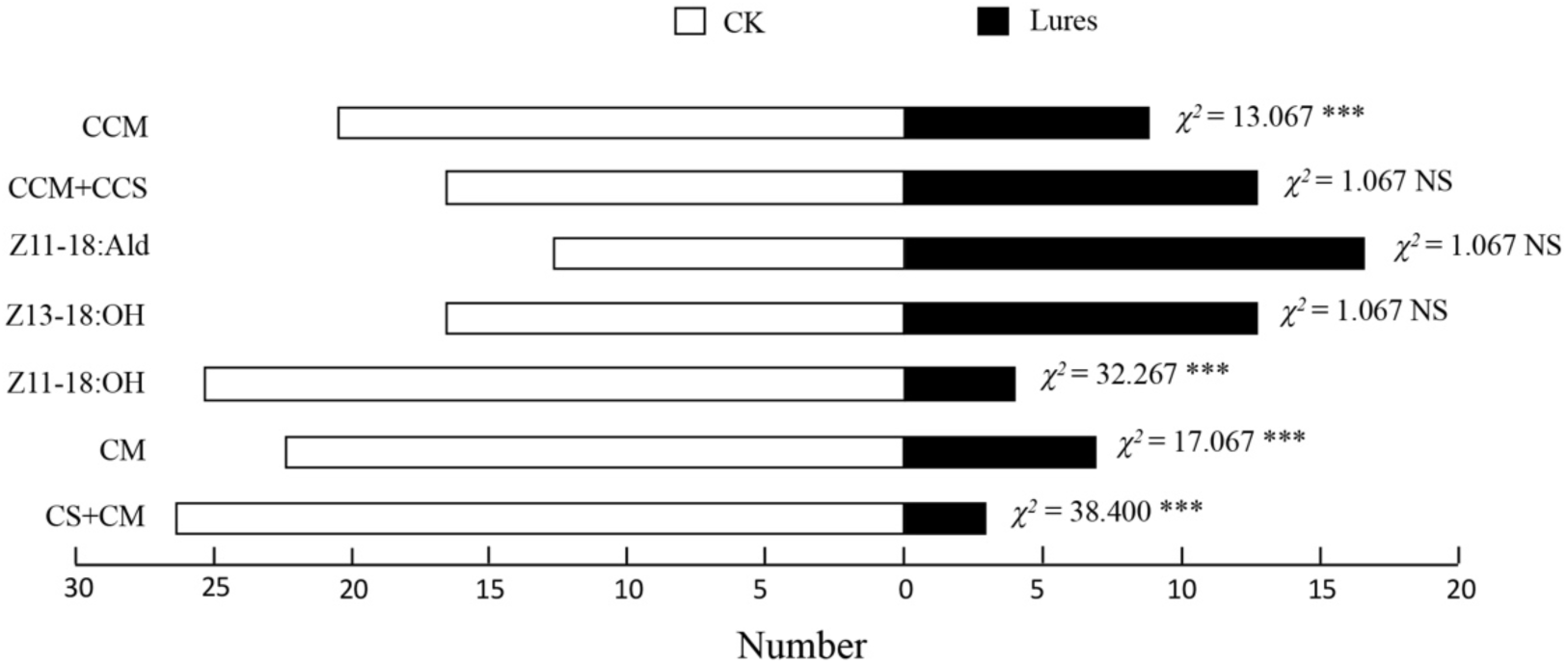
Response of male *C. suppressalis* to seven lures over controls in an olfactometer. CCM means commercial *C. medinalis* sex pheromone; CCM+CCS means combination of commercial *C. medinalis* and *C. suppressalis* sex pheromones; CM means the four components of *C. medinalis* sex pheromone (Z13-18:Ald, Z13-18:OH, Z11-18:Ald, Z11-18:OH=100:36:11:24); CS+CM means combination of the components of *C. suppressalis* and *C. medinalis* sex pheromones (Z11-16:Ald, Z13-18:Ald, Z9-16:Ald, Z13-18:OH, Z11-18:Ald, Z11-18:OH=46:106:5:36:11:24). The concentrations of the lures are 1 μg/μl except the two commercial sex pheromones. The control (CK) is redistilled hexane. Significance levels of χ^2^ (*Chi*-square test) indicated by *** (*P* < 0.001) or NS (no significant difference). *N* = 30.

### Adding Z11-18:OH and Z13-18:OH to the CCS significantly decreased the captures of *C. suppressalis* males in the field experiment

We further determined the inhibition effects of sex pheromone components of *C. medinalis* on response of *C. suppressalis* male in the field. The results showed that adding Z11-18:OH (*t* _(26)_ = 2.334, *P* < 0.05) or Z13-18:OH (*t* _(26)_ = 2.252, *P* < 0.05) to the CCS obviously inhibited the capture of *C. suppressalis* male in the field (Fig. 4). However, compared with control treatment (CCS), adding Z11-18:OH (*t* _(26)_ = 2.042, *P* = 0.051) has no significant difference on response of *C. suppressalis* male. Which suggested that adding Z11-18:OH or Z13-18:OH to the CCS could significantly inhibit the attraction of CCS to *C. suppressalis* male in the field.

**Fig. 4.**
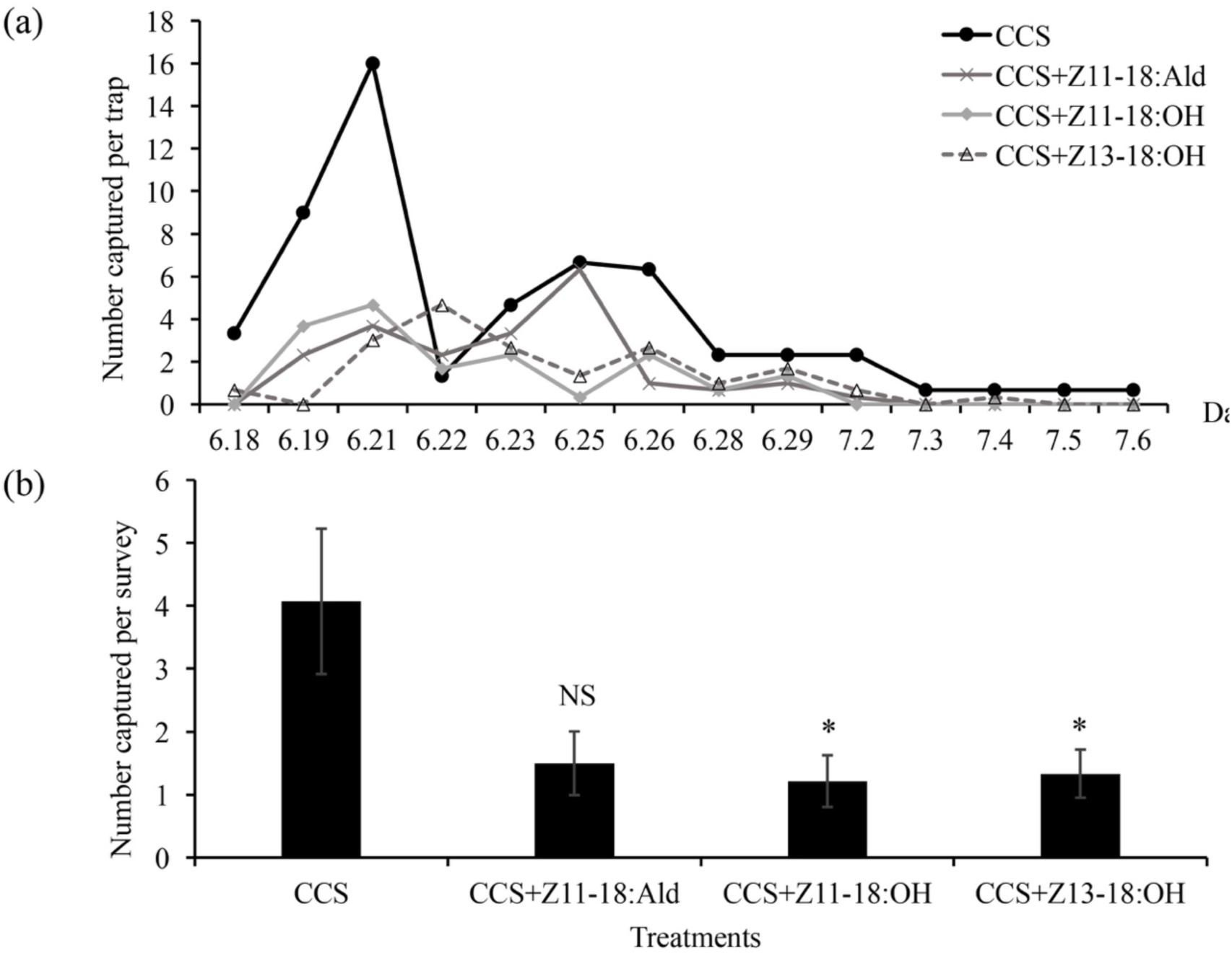
The dynamic linear graph (a) and summary graph (b) of number of *C. suppressalis* males captured by commercial *C. suppressalis* sex pheromone alone (CCS) and combined with the components of *C. medinalis* sex pheromone (Z11-18:Ald, Z11-18:OH, Z13-18:OH) in Jiangxi, China, from June to July 2014. The difference of capture of *C. suppressalis* males between treatments and control treatment were analyzed with a *t*-test. *, *P* < 0.05; NS, no significant difference.

We further investigated the response of combination Z11-18:OH, Z13-18:OH and CCS to the *C. suppressalis* male. We also found that adding both Z11-18:OH and Z13-18:OH (*t* _(24)_ = 4.677, *P* < 0.001) to the CCS could significantly inhibit the attraction of CCS to *C. suppressalis* male in the field (Fig. 5). The means of total captures by CCS without and with both Z11-18:OH and Z13-18:OH were 18.66 and 2.33, respectively.

**Fig. 5.**
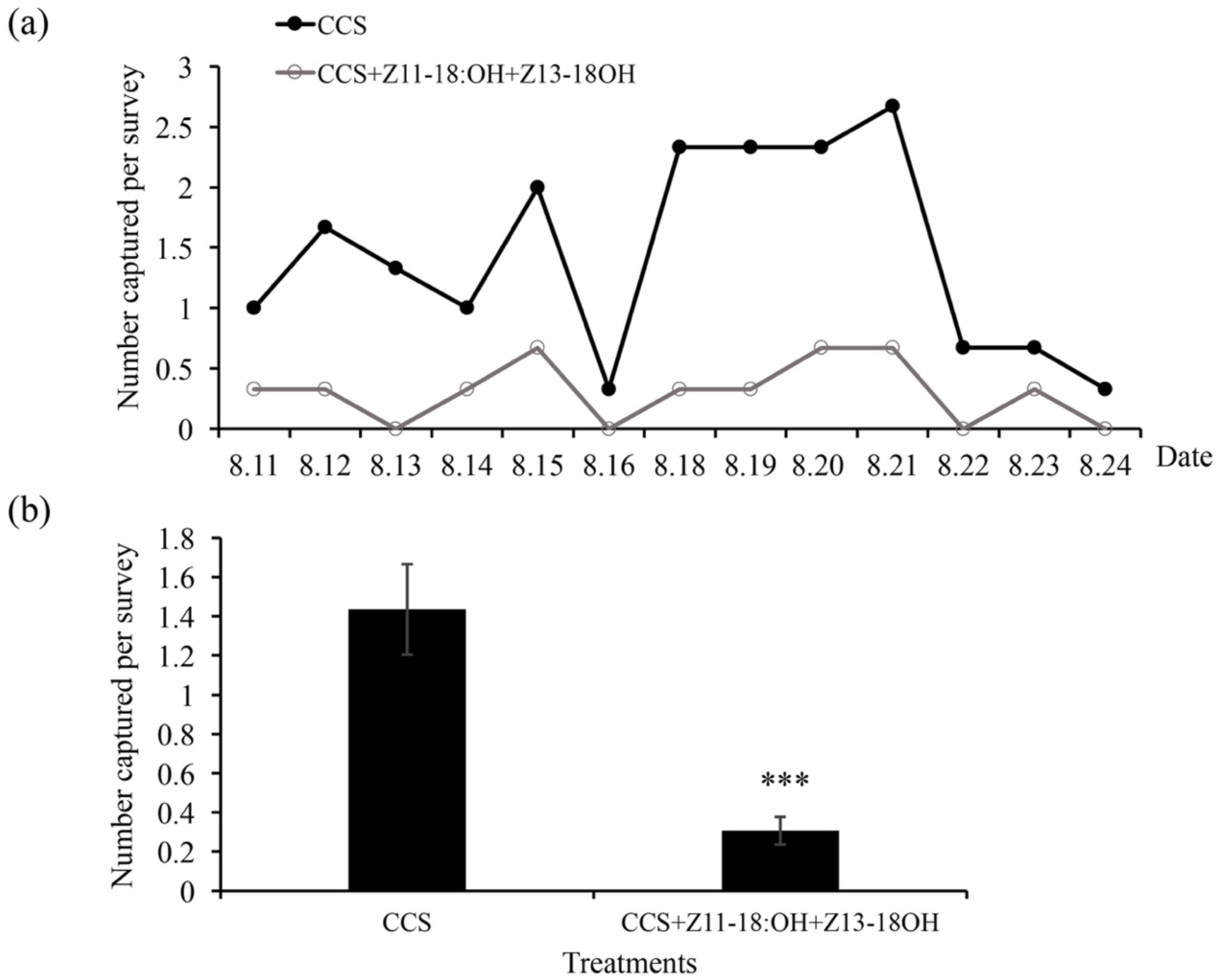
The dynamic linear graph (a) and summary graph (b) of number of *C. suppressalis* males captured by commercial *C. suppressalis* sex pheromone alone (CCS) and CCS plus individual components (Z11-18:OH and Z13-18:OH) in Jiangxi, China, from 11th to 24th August, 2014. The difference of capture of *C. suppressalis* males between treatments and control treatment were analyzed with a *t*-test. ***, *P* < 0.001.

## Discussion

In this study, we found that mixing the sex pheromones of *C. suppressalis* and *C. medinalis* interfered the capture of *C. suppressalis* male in the field experiment. Our finding agreed with the result of Gemeno et al. (2006), which has shown that mixing sex pheromones of two insect species resulted in significantly lower captures of one of these species. Similarly, attraction of *Tetanolita mynesalis* (Lepidoptera: Noctuidae) to its own pheromone was inhibited when mixing with the sex pheromone of *Lacinipolia renigera* (Lepidoptera: Noctuidae) (Haynes et al. 2002), and the sex pheromone of *Adoxophyes orana* (Lepidoptera: Corynidae) inhibited the attraction of *Cydia pomonella* (Lepidoptera: Tortricidae) (Potting et al. 1999). Such phenomena of inhibition or antagonism in the sex pheromone systems of several insect species probably contribute to the reproductive isolation of closely related species (same genus) (Roelofs and Cardé 1974, Borden 1997, Cardé and Haynes 2004). In addition, pheromone inhibition or antagonism may occur between species that are not closely related (different genera). For instance, Lopez et al. (1990) found that multispecies sex pheromone traps caused a reduction of baiting in more than one species that the baiting in individual species traps. Our results demonstrated that CCM inhibited the attraction of *C. suppressalis* and we did not recommend the use of these two sex pheromones together in the field.

Sex pheromones in many insect species are composed of multiple components. In such cases, air permeation with an individual component can often disrupt the sexual communication between male and female. Our olfactory experiments and field tests showed that adding Z11-18:OH and Z13-18:OH to the traps baited with CCS caused the antagonism in attraction of *C. suppressalis* male. The results are similar to the studies of the yellow stem borer *Scirpophaga incertulas* (Lepidoptera: Pyralidae), which showed that Z-11-hexadecenol from *S. incertulas* sex pheromone inhibited the captures of *C. suppressalis* males (Cork and Basu 1996). Some alcohols have been shown could be synergists or inhibitors of many insect pheromones. Yu et al. (2014) reported that 1-undecanol acted as sex pheromone synergist to enhance the attraction of male *Grapholita molesta* (Lepidoptera: Tortricidae) pheromone traps. Several previous studies have demonstrated that Z-11-hexadecenol inhibited the attraction of male *Mamestra brassicae* (Lepidoptera: Noctuidae) (Struble et al. 1980), as well as the attraction of *Cydia pomonella* (Lepidoptera: Tortricidae) was inhibited by Z-9-tetradecenol (Chisholm et al. 1983). Our results indicated that the alcohols Z11-18:OH and Z13-18:OH may act as the inhibitors for *C. suppressalis* sex pheromone.

In summary, we found that combined using of CCS and CCM in the trip caused no capture of *C. suppressalis* male in the field experiment. Our results also demonstrated that Z11-18:OH and Z13-18:OH could improve the efficiency of mating disruption by inhibiting the attraction of male *C. suppressalis* to CCS. Therefore, we do not suggest that using sex pheromones of these two species together in the field. Our finding is very important to guide farmers in agricultural activates. In addition, we also suggest that Z11-18:OH and Z13-18:OH could be potential repellents or antagonists of *C. suppressalis* sex pheromone, which may contribute to biological control in sex pheromone-mediate mating disruption in future.

## Acknowledgments

The authors are grateful to Zhiwen Yao for assistance in arthropod sampling. This research is financially granted by the National Key Research and Development Program of China (Project No. 2017YFD0301604, 2016YFD0200808 and 2017YFD0200400). M.L. acknowledges the support from the China Scholarship Council (CSC) (File No. 201708360095).

